# Imaging mechanical properties of sub-micron ECM in live zebrafish using Brillouin microscopy

**DOI:** 10.1101/491803

**Authors:** Carlo Bevilacqua, Hector Sanchez Iranzo, Dmitry Richter, Alba Diz-Muñoz, Robert Prevedel

## Abstract

In this work, we quantify the mechanical properties of extra-cellular matrix (ECM) in live zebrafish using Brillouin microscopy. Optimization of the imaging conditions and parameters, combined with careful spectral analysis, allows us to resolve the thin ECM and distinguish its Brillouin frequency shift from the surrounding tissue. High-resolution mechanical mapping further enables to directly measure the thickness of the ECM label-free and *in-vivo*. We find the ECM to be ~400nm thick, in excellent agreement with electron microscopy quantification. Our results open the door for future studies that aim to investigate the role of ECM mechanics for zebrafish morphogenesis and axis elongation.

## 1. Introduction

At the heart of morphogenesis lies a tight coordination between specific patterns in mechanical forces, generated by the cytoskeleton and the extracellular matrix (ECM), and patterns of signalling proteins that instruct and modify the cytoskeleton and the cell’s fate. Such systems are intrinsically coupled in mechanochemical feedback loops, exemplified by work showing how ECM stiffness can drive cell fate [1]. In textbook examples of organ elongation, such as germband extension in Drosophila or neural plate extension in vertebrates, the force anisotropies that instruct shape are generated within the cells. In comparison, how the mechanical properties of the ECM influence organ elongation is less well understood. Specifically challenging has been the study of the formation, elongation and branching morphogenesis of tubes. Such morphological events are crucial for the development and function of organs such as the mammary ducts, the kidney tubules or lung alveoli. Several key cellular activities can drive tubule elongation [2]. Moreover, Crest *et al.* recently showed how patterning anisotropic resistance within the ECM can drive *Drosophila* follicle elongation [3]. Determining to what extent such mechanics are conserved in vertebrates would require ECM stiffness measurements in 3D.

Measuring these mechanical forces and properties at high spatial resolution in live, threedimensional tissues is challenging [4,5]. Among the approaches to measure micrometer-sized mechanical properties of live cells and tissues, such as elasticity or viscosity, Atomic Force Microcopy [6] (AFM) and micropipette aspiration [7] are most often used. However, both require the transmittance of contact forces which could perturb the sample, are restricted to cell surfaces only, and rely on mechanical models to extract local elasticity parameters. Optical approaches such as magnetic twisting cytometry [8] or optical tweezers [9] measure viscoelastic properties from cellular deformations induced by either magnetic fields or high power laser beams, which also limit their application to live tissues. Recently, methods have been put forward that involve the injection of micro-beads or lipid-coated droplets into 3D tissues or large cells to actively apply forces to them with optical tweezers or magnetic fields [4,10]. While such approaches provide elasticity information in an *in-situ* context, the invasive introduction of foreign matter into live biological samples is likely to interfere with sensitive biological processes such as those encountered during embryo development and only provide limited spatial information.

Brillouin microscopy has recently emerged [11] as a new technique in the field of mechanobiology to explore the mechanics of living cells and tissues. Brillouin microscopy is an all-optical method that probes the visco-elastic properties of a material via light scattering. Thus, it achieves high spatial, i.e. near diffraction-limited resolution in 3D. Furthermore, it does not require artificial labelling nor external loading of the sample, thus making it a promising label-free and non-contact method to study mechanical properties of living tissues. In recent years, Brillouin microscopy has enabled applications in cell biology [12–14], ophthalmology [15,16], as well as disease detection [17]. Previous works have established that it is in principle feasible to measure cell mechanical properties with subcellular resolution [12,13]. Furthermore, Schlüssler *et al.* have used Brillouin microscopy for the biomechanical assessment of regenerating tissues in living zebrafish [18] and Palombo *et al.* have studied the contribution to the Brillouin shift of different ECMs from various rat fibrous connective tissue *ex-vivo* as well as their relative fiber alignment [19]. Lastly, Elsayad *et al.* showed how Brillouin microscopy could be used to map the viscoelastic signature of the ECM of *Arabidopsis thaliana* and their relation to the hydrostatic pressure of the cells within [20].

Here, we aim to quantify stiffness of thin ECM layers *in-vivo*, thereby overcoming some of the limitations present in those previous studies. Specifically, we focus on a vertebrate tissue that undergoes elongation, the zebrafish notochord. The notochord is an embryonic midline structure common to all members of the phylum Chordata that gives rise to the vertebrae and contributes to the center of the intervertebral discs [21]. It provides structural support to the developing embryo and secretes inducible signals to several tissues. In zebrafish, the notochord is formed by two cell types: sheath and vacuolated cells. Sheath cells are epithelial cells that form a tube-like structure and basally secrete ECM, while vacuolated cells are located apically in the lumen and provide hydrostatic pressure to straighten the antero-posterior body axis of the embryo [22,23]. The ECM is formed by three layers: a thin inner laminin-rich basement membrane layer [24], a middle layer of densely packed collagen fibers (especially rich in collagen type-II which is also a major constituent of cartilage [21]), and an outer layer in which extracellular fibers run perpendicularly to the middle layer [25]. If too many vacuolated cells die in the same embryo, for example in *cavin1*-knockout embryos, the resulting fish is unable to sustain the necessary intra notochord pressure and will have a shorter antero-posterior axis [26]. Moreover, a loss in ECM integrity (by for example inhibiting cross-linkers of collagen [27]) also leads to a shorter and kinked axis, highlighting how both vacuolated and sheath cells are crucial for notochord function. Being able to measure the mechanical properties of ECM *in-vivo* during development could open the door to study its role in morphogenesis and organogenesis.

## 2. Experimental methods

### Brillouin scattering

Brillouin microscopy uses visible or infrared monochromatic (laser) light to probe the mechanics of a material through light scattering from thermally induced, gigahertz-frequency acoustic modes, i.e. ‘phonons’. A typical spectrum from Brillouin scattering will consist of a symmetric pair of peaks – the Brillouin Stokes and anti-Stokes peaks – on each side of the elastically scattered (Rayleigh) input light, which is many orders of magnitude more intense. This spectrum provides information about the local elasticity of the sample. Specifically, the change in frequency (GHz) of the Brillouin scattered light due to photon-phonon interactions is given by:

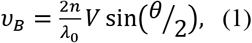

where *n* is the material refractive index within the scattering volume, λ_0_ is the incident wavelength, θ is the angle between the incident and scattered light, and *V* is the medium’s sound velocity. The acoustic velocity *V* is then related to the complex longitudinal modulus, *M*, of which the real part, defined as the ratio of stress to strain in a uniaxial strain state, is given by *M’* = *ρV*^2^, where *ρ* is the density. The linewidth **Δ** (full-width at half-maximum, FWHM) of the Brillouin spectrum is related to the imaginary component of M, i.e. the loss modulus, which is a measure for viscosity, by *M”* = *ρV*^2^Δ/*υ*_*B*_. Mapping frequency shifts and linewidth in space across a specimen will thus lead to a 3D resolved mechanical image.

Typically, the spontaneous Brillouin scattering signal is weak, which has in the past required the use of high laser powers and long acquisition times [28], which however are often incompatible with meaningful applications in biology. In the last decade, the development of alternative spectrometer designs [11], based on the virtually-imaged phased array (VIPA), has greatly increased the spectral detection efficiency thus enabling mechanical characterization of biological tissue at safe power levels and opening up new avenues for *in-vitro* and *in-vivo* studies.

While Eq. 1 requires knowledge of *ρ* and *n*, i.e. the density and refractive index to compute an absolute elastic modulus, previous studies have shown that the variation of the ratio ρ/ n^2^ is fairly small for various biological samples [12,29]. Therefore, the Brillouin shift itself is a good approximation for the longitudinal modulus. In general, the longitudinal modulus in biological material (e.g. tissues, cells, biopolymers) is several orders of magnitude higher than the traditional quasi-static Young’s modulus, due to the incompressibility of the material and the frequency dependence of the modulus. The relationship between the high-frequency longitudinal modulus probed by Brillouin scattering and the traditional quasi-static Young’s modulus which is often used in the field of mechanobiology has yet to be fully understood for biological tissues. In our study, we report the Brillouin shift as a metric of mechanical properties of the zebrafish embryos as it is the direct parameter measured in the experiment.

### Imaging setup

Our confocal Brillouin microscope (Fig. 1) is conceptually based on a two-stage VIPA spectrometer design adapted from Scarcelli *et. al.* [30]. The laser source is a single-frequency mode, 532-nm continuous-wave (CW) laser (Torus, Laser Quantum), that is single-mode coupled to ensure a Gaussian beam profile. After collimation, the illumination light passes a half-wave plate (HWP) and polarized beam splitter (PBS), which serve to control the intensity of the light being sent to the microscope and calibration arm, respectively. Afterwards, the beam enters the microscope body (Zeiss Axiovert 200M) through a sideport and is then focused into the sample by an objective lens (either Zeiss Plan-Apochromat 40x, 1.0NA with adjustable numerical aperture (NA) or Zeiss Plan-Apochromat 40x 1.4NA). This allows us to vary the extent of the optical point-spread function (PSF) and therefore the volume over which the Brillouin scattered light is captured (Fig. 1B). The microscope is further equipped with a Piezo translation stage (P-545.3R8H, Physik Instrumente) that allows positioning of the sample with nanometer precision.

**Fig.1:**
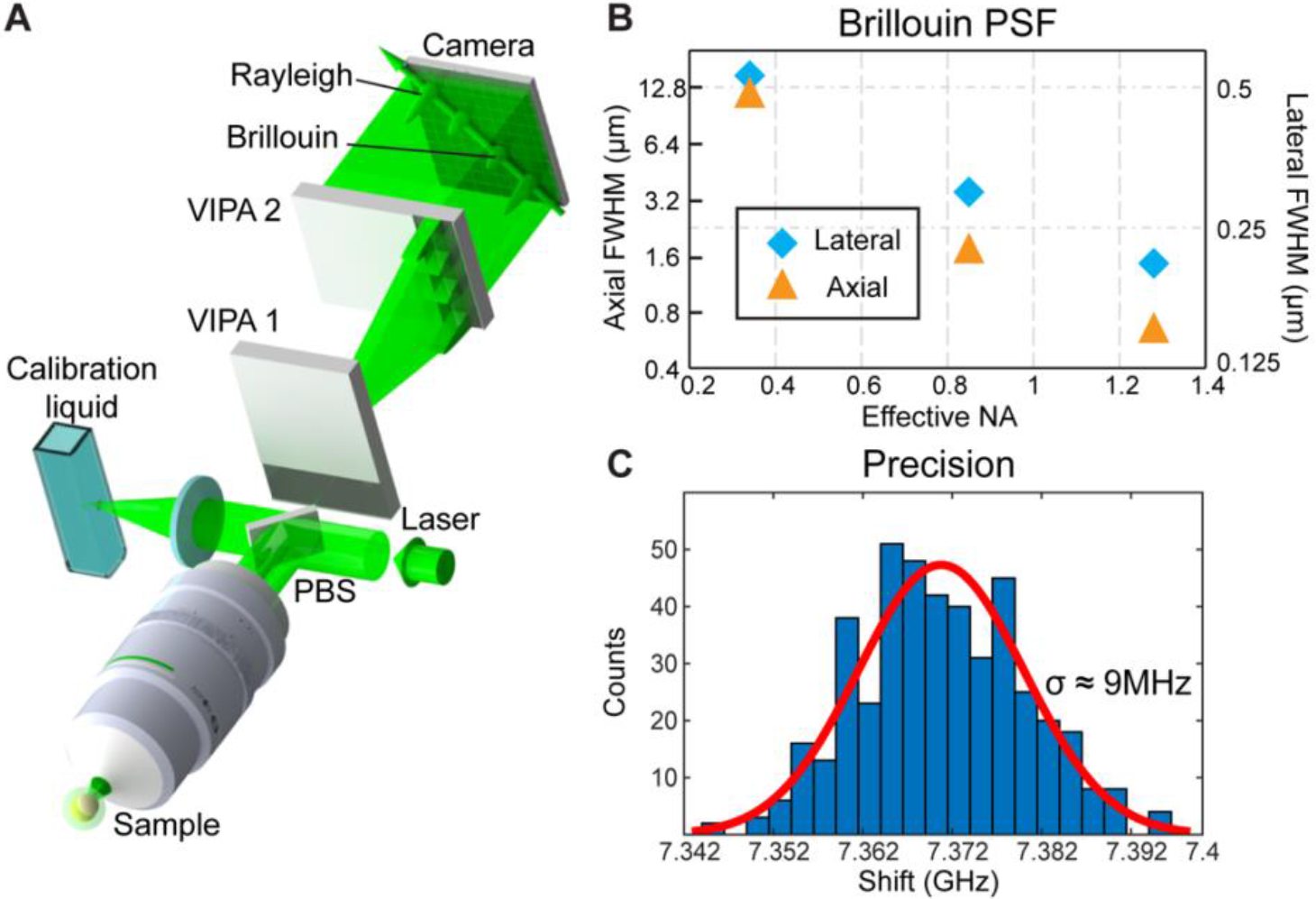
Brillouin microscope setup and characterization. (A) Schematic of the dual-stage VIPA based imaging setup. PBS - polarizing beamsplitter. (B) Experimentally recorded PSF width as a function of effective objective NA (linear-log scale). This was measured by scanning the focal volume across an oil/glass interface and fitting an Erf-function to the obtained edge response of the amplitude of the Brillouin spectrum of oil. This also determined the effective NA. (C) Brillouin shift precision, obtained from 440 individual measurement of water under the same conditions used *in-vivo* (8.3 mW, 180 ms camera integration time).

The Brillouin scattered light is collected in backscattering direction by the same objective and sent to the Brillouin spectrometer by coupling into a single-mode fiber, which also ensures confocality of the light detection. A quarter-wave plate (QWP) before the PBS ensured that most of the backscattered light collected by the objective lens is delivered to the spectrometer. The Brillouin spectrometer consists of a two-stage virtually imaged phased array in cross-axis configuration, similar to a previous report [30], but with the addition of a Lyot stop [31] before the camera (iXon DU897, EMCCD camera; Andor Technology). The spectrometer features a free spectral range of 30GHz, with a nominal spectral resolution of 460MHz (finesse ~65). The VIPA cross-axis and Lyot stop configuration yield a high, ~65dB elastic background suppression, sufficient for *in-vivo* imaging of the zebrafish embryo.

Our confocal Brillouin microscope furthermore offers the ability to perform simultaneous or sequential confocal fluorescence imaging, by employing the Brillouin or a separate CW 488nm laser, respectively. The fluorescence light is separated from the Brillouin light by a narrowband bandpass filter (Semrock LL01-532), and detected by a proper combination of a focusing lens (f=75mm), a pinhole (40μm) and a photomultiplier tube (Thorlabs PMT1001).

The acquired scattering spectra are analyzed in real time with custom-written Labview program by fitting Lorentzian functions to obtain the position, width and intensity of the Brillouin peaks. In order to calibrate the frequency axis, a reference measurement of a water sample is acquired after every 50 measurements (~10s). The conversion between camera pixels and GHz is performed assuming a linear relationship between the distance of the Brillouin peaks on the camera (*d*_*p*_) and the corresponding Brillouin shift (*s*): *s*=(*FSR-*α*·d*_*p*_)/2, where the coefficient *α* (GHz/pixel) is measured from the calibration spectrum of water (shift of 7.4GHz at room temperature). The water sample is placed in the transmitted beam path of the PBS (c.f Fig.lA) that acts as a separate calibration arm, and mechanical shutters switch between the lightpaths. This recalibration compensates for omnipresent frequency drifts in the sub-GHz regime of the laser, and specifically to correct the drift of the spectral axis on the camera and to update the conversion coefficient *α* during the experiment. Altogether, this ensures a high, ~9MHz, precision in our frequency shift measurements under *in-vivo* experimental conditions (Fig. 1C).

### Zebrafish experiments

For all Brillouin and electron microscopy (EM) images a Tg(*col9a2*:GFPCaaX)^pd1151^ zebrafish line [26] that labels sheath cells was used. For illustration purposes in Figure 2 a triple transgenic Tg(*col9a2*:GFPCaaX)^pd1151^; SAGFF214A [32]; Tg(UAS-*E1b*:NfsB-mCherry)^c264^ [33] labelling sheath and vacuolated cells was used. In all experiments, 1-phenyl 2-thiourea (PTU) was added at 0.003% concentration at 10 hours post fertilization to avoid pigmentation. For Brillouin imaging, fish were immobilized on one of their sides in 1% agarose with 0.003% PTU and 0.016% tricaine. After the Brillouin imaging experiments, the larvae were released from the agarose and behaved normally after cessation of sedation.

**Fig.2:**
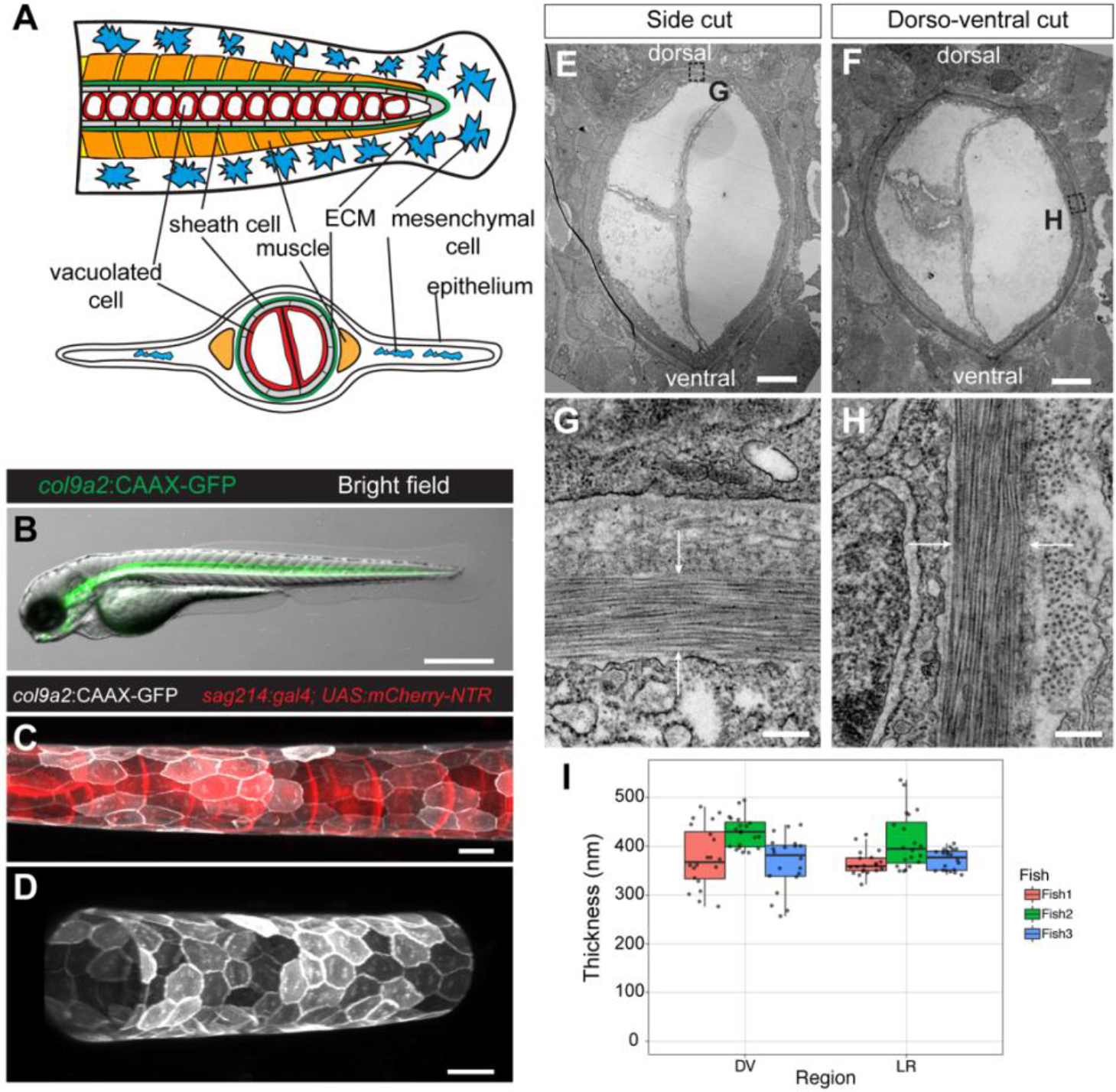
Characterization of the zebrafish notochord at 3 days post fertilization (dpf). (A) Schematic of the zebrafish tail at 3 dpf. Lateral (top) and transversal (bottom) views. (B) Bright field with overlapping fluorescence of a Tg(*col9a2*:GFPCaaX) zebrafish embryo at 3 dpf with GFP fluorescence from sheath cells. (C-D) Maximum intensity projection of the notochord in triple transgenic Tg(*col9a2*:GFPCaaX); SAGFF214A; Tg(UAS-*E7b*:NfsB-mCherry) zebrafish embryo at 3 dpf. Both sheath and vacuolated (C) or only sheath cells (D) can be observed. (E-F) Low magnification TEM of transversal notochord after side (E) and dorso-ventral (F) cuts of a Tg(*col9a2*:GFPCaaX) zebrafish embryo at 3 dpf. (G-H) High magnification TEM of transversal view ECM (between white arrows) after side (G) and dorso-ventral (H) cuts of a Tg(*col9a2*:GFPCaaX) zebrafish embryo at 3 dpf. (I) Quantification of ECM thickness in left-right (LR) and dorso-ventral (DV) regions of 3 embryos. Scale bars, 500μm in (B), 20μm in (C and D), 5μm in (E and F) and 250nm in (G and H).

For EM imaging, samples were chemically fixed by immersing them in 2.5% glutaraldehyde and 4% paraformaldehyde in 0.1M PHEM buffer. Sections were post-stained with uranyl acetate for 5 minutes and with lead citrate for 2 minutes. The overall EM protocol is similar to Ref. [34,35].

### 3. Results

After assessing the notochord geometry by imaging available zebrafish lines that mark sheath and vacuolated cells (Fig. 2A-D), and in order to assess the ultrastructure of the zebrafish ECM, we performed transmission electron microscopy (JEOL 2100 at 120kV) on 70nm thick crosssections collected from 200-250μm before the end of the notochord. We performed both side and dorso-ventral cuts to avoid compression artefacts due to the pressure of the diamond knife on the sample and measured the ECM thickness across 20 locations in each fish in the region of the notochord perpendicular to the cut (Fig. 2E-H). We observed that even with embryo-to-embryo variability the thickness of the medial ECM layer is consistent around the notochord in all the samples measured, with a dorso-ventral average of 390 ± 58 nm and a left-right average of 383 ± 43 nm (mean ± S.D.) (Fig. 2I).

Next, we set out to explore to which extent we could resolve the ECM *in-vivo* using Brillouin microscopy (Fig. 3). To do so, we first acquired an overview image with a 0.85NA objective over a ~200×200μm field-of-view (FOV) centred approximately 500μm anterior from the posterior end of the notochord and across the approximate middle plane of the lateral axis (Fig. 3B). Plotting the Brillouin frequency shift, the thin structure of the ECM becomes indeed visible due to the expected high Brillouin shift of the ECM’s collagen fibers [19], even when spatially undersampling the image acquisition (1μm step size compared to ~400nm ECM thickness). We further confirmed that the ECM was indeed the source of the high Brillouin shift by comparing the Brillouin image with a subsequently recorded confocal fluorescence image at the same focal position, as the ECM secreting sheath cells where labelled with GFP (Fig. 3A). Based on these recordings, we performed further Brillouin imaging at high spatial sampling (100nm step size) over a ~5×5μm region (Fig. 3C,D) centred 250μm anterior from the posterior end of the notochord, which indeed identified the ECM as a 400-500nm thick region with relatively high Brillouin shift. We further observed that the Brillouin spectrum in the central region displayed a strong asymmetry towards higher Brillouin shift, indicative of a second spectral peak. By fitting the spectrum with the sum of two independent Lorentzian functions, we can indeed separate the ECM and surrounding tissue components and their respective contribution to the overall Brillouin spectrum (Fig. 3E). Compared to previous work [36] which performed similar analysis by subtraction the cell’s buffer medium from the overall spectrum, our direct fitting approach also works in heterogeneous tissues and is less susceptible to (laser drift induced) artefacts. Thus, we find that careful spectral analysis allows distinguishing mechanical properties of different mechanical components even in a regime where the structure size approaches the optical resolution of the Brillouin microscope. In particular, we find the ECM to display a significantly higher shift of 9.98 ± 0.10 GHz (*n*=5 fish) when averaged along the ECM and across individual fish, significantly higher than the surrounding cells (7.86 ± 0.02 GHz). This is in good agreement with *ex-vivo* studies of collagen type-II in rat cartilage, performed with a high spectral resolution Fabry-Perot spectrometer [19]. Furthermore, analyzing the Brillouin spectrum across the ECM allows us to characterize the ECM thickness *in-vivo* and in a label-free manner by plotting the ratio of the respective amplitudes of the mechanical signals (Fig. 3E-G). The amplitude of the ECM contribution to the overall spectrum is a convolution of the optical PSF and the physical size of the ECM (approximated by a box-like function). Since we have characterized the Brillouin PSF of our microscope (Fig. 1B), we can thus estimate the physical thickness of the ECM by deconvolution of the measured line response. This yielded an average ECM thickness of 380 ± 78 nm (*n*=5 fish), in excellent agreement with the EM image analysis (Fig. 2I). This shows that with diligent spectral analysis it is indeed possible to characterize the mechanical properties of thin ECM in living organisms.

**Fig.3:**
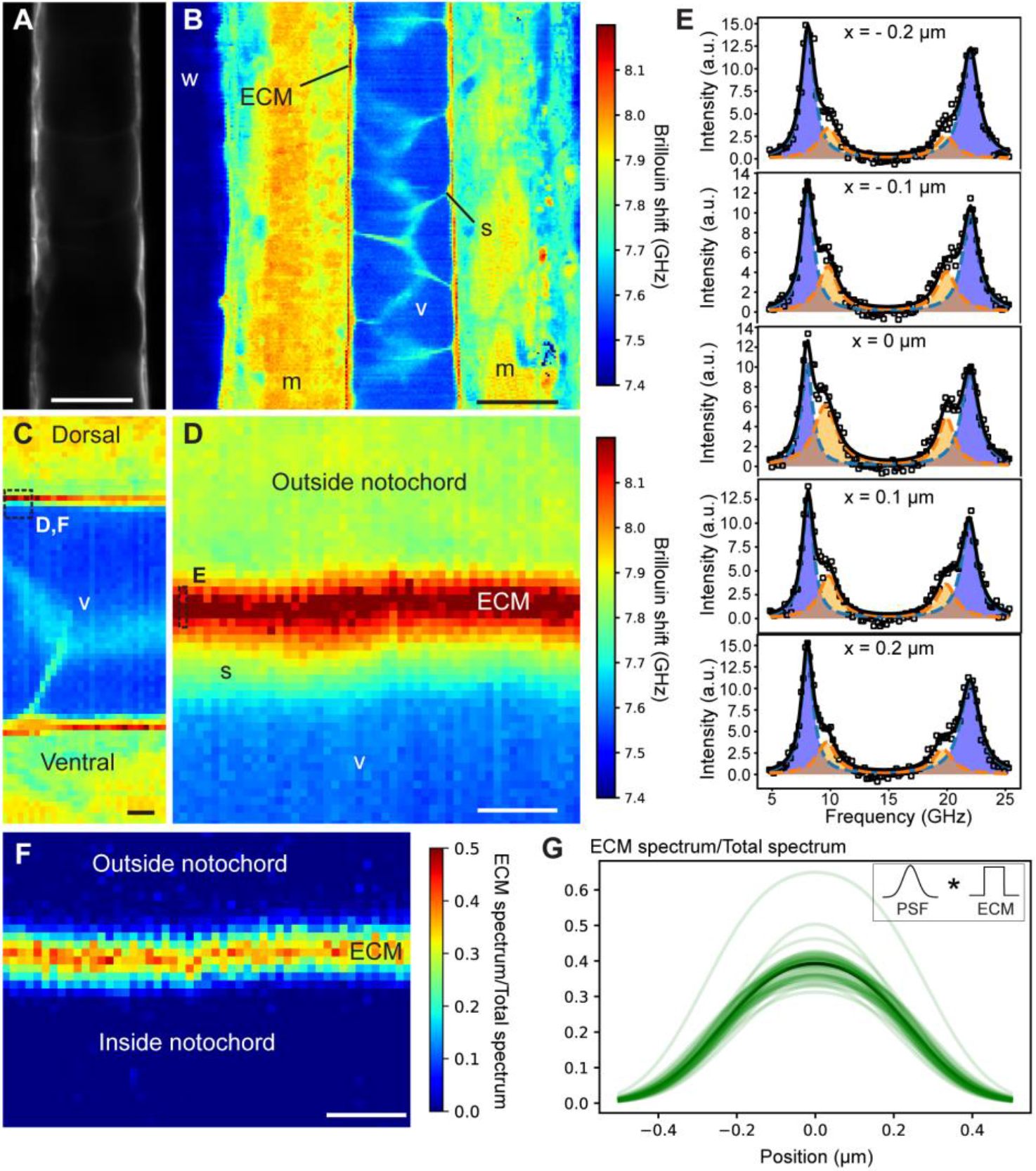
High-resolution mechanical imaging of zebrafish notochord ECM *in-vivo*. Brillouin shift maps located 200-250μm before the end of the notochord, and approximately in the middle between the left-right animal’s side. (A) Confocal image of sheath cells (GFP) as well as corresponding coarse-grained overview Brillouin image in (B) recorded with 1μm step size at 3 days post fertilization (dpf). (C) Zoom-in of Brillouin shift map at 3 dpf. (D) Brillouin map of boxed area denoted in (C) with a step size of 0.1μm. (E) Brillouin spectra recorded at discrete positions across the ECM (boxed area in (D)). Double-peak fitting distinguishes the Brillouin shift of the ECM (orange) and surrounding tissue (sheath cells and outside notochord - blue). Here, the fitting parameters were seeded from the parameters obtained from the fit to the pixel with highest ECM peak contribution and then subsequently constraining the ECM peak around ± 0.7GHz when analyzing the surrounding pixels in order to prevent artefacts. We note that the observed variability in ECM peak shift is much lower than this fitting constraint (see main text). A spatial map of the ratio of the spectral contributions is plotted in (F). (G) Line plots of ECM contribution to total spectrum across the ECM in (F), centered by their respective maximum. Inset shows convolution of measured line response. Brillouin images were obtained with a 0.85NA objective and using 8.3mW of laser power and 0.25s of exposure time per pixel. w, water; m, muscle; s, sheath cell; v, vacuole, ECM, extracellular matrix. Scale bars, 20μm in (A,B), 5μm in (C), 1μm in (E,F).

In order to assess the ECM in three-dimensions, we also performed axial imaging of the zebrafish (Fig. 4). Here we again observed that the Brillouin shift map obtained from a single-Lorentzian peak fit hardly revealed the ECM, and that the contribution of the ECM was only visible in a narrow, 10-20μm region surrounding the dorso-ventral midline (Fig. 4A). Also, the area of high shift (> 8.1 GHz) showed a strong dependence on the effective NA used for the Brillouin imaging (Fig. 4 A,E,I). In particular, we found that a relatively medium NA of 0.85 yielded the best contrast. Overall, a double-peak fit analysis again led to a higher contrast of ECM when compared to the surrounding tissue (Fig. 4B-D,F-H,J-L). Here, the region over which the ECM had a clear contrast could be extended to up to 50μm in the axial direction. Finally, we note that when the PSF was fully overlapping with the ECM (Fig. 4M, right panel) the ‘tissue’ peak displayed a slight, ~200MHz shift to higher frequencies compared to regions outside the ECM. Since in this condition we don’t expect any major contribution to the spectrum from the surrounding cells, we attribute this to the observation of two acoustic modes (bulk and parallel-to-surface) within the ECM, consistent with recent experiments by Palombo *et al.* [19].

**Fig.4:**
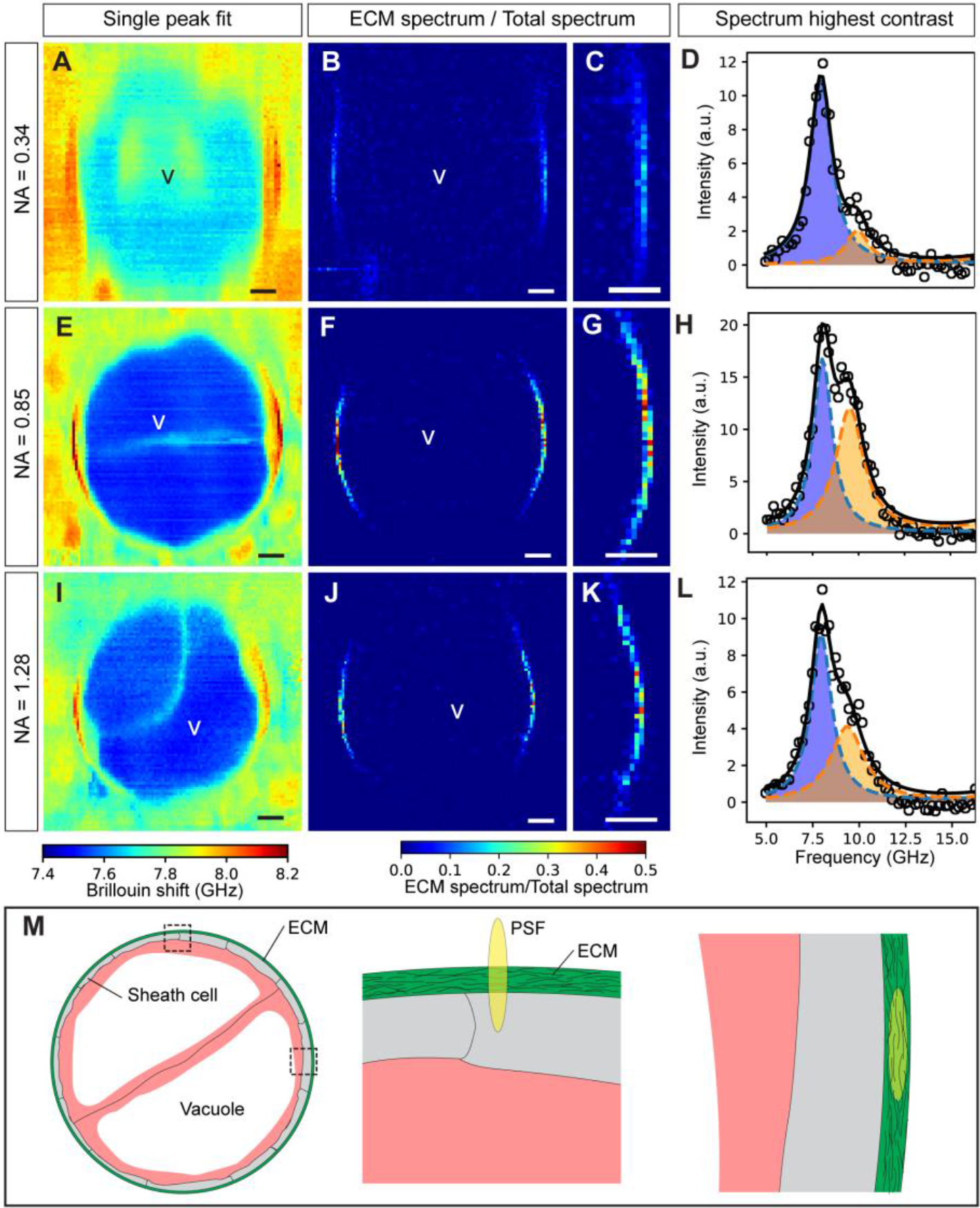
Axial cross-section of zebrafish notochord mechanics. The effect of objective NA on the Brillouin map and contrast of the ECM is shown for low (0.34), medium (0.85) and high (1.28) effective NA in A-D, E-H and I-L respectively. In traditional single-peak fit Brillouin shift maps the ECM displays poor visibility (A,E,I). Dual-peak analysis and plotting the spectral ratios does improve contrast (B,F,J). (C,G,K) Zoom-ins. (D,H,L) Spectra of pixels with the highest ECM spectrum / Total spectrum contrast in (C,G,K), respectively. (M) Illustration showing the crosssection of the notochord (left) as well as the overlap of the PSF with the ECM when measuring on the lateral (middle) and dorso-ventral (right) side, respectively. Brillouin images were obtained using 8.3mW of laser power and 0.25s of exposure time per pixel. In panels A, B, C, E, F, G, I J, K dorsal is located to the left and ventral to the right. Step size: 0.5μm. v, vacuolated cells. Scale bars, 10μm.

### 4. Discussion and outlook

In this work, we showed for the first time that Brillouin microscopy is capable of measuring the mechanical properties of the ECM *in-vivo* and with high spatial resolution inside a vertebrate organism. This was achieved by imaging system optimization and careful spectral analysis. In particular, diligent multi-peak spectral fitting allowed to clearly separate the mechanical properties of the ECM from the surrounding tissue, which so far was only demonstrated with Fabry-Perot based Brillouin spectrometers that typically feature much higher spectral resolution (at the expense of lower efficiency and imaging speed typically prohibitive of *in-vivo* measurements). Furthermore, our high-resolution mechanical measurements were able to determine the physical size of the ECM, in excellent agreement with ultrastructural analysis based on EM images.

The fact that we have achieved the optimal ECM Brillouin signal contrast with the use of a medium NA objective might appear counter-intuitive at first sight. However, one has to consider the spatial overlap of the axially elongated PSF with the structure of the ECM (Fig 4M). Given the ECM thickness, geometry and radius, a slightly elongated PSF (~0.3 and 1.8μm in x,y and z-direction, respectively for our 0.85NA objective) has the optimal volumetric overlap and is thus preferable in this case. Another point worth discussing is the observation that in Fig. 4 the ECM is not visible on the lateral sides. We attribute this largely to the small PSF-ECM overlap in combination with the dependence of the longitudinal modulus on the ECM collagen fibers’ orientation, as previously characterized in rat ECMs [19]. In particular, when probing the fibers in an orthogonal (isostress) direction, as is the case on the lateral sides of the animal given our microscope arrangement, the spectral shift is expected to decrease by 10-15% [19].

Finally, one has also to consider the fact that if the axial extent of the PSF becomes smaller than the attenuation length of the acoustic modes, only an average mechanical property will be probed, i.e. one would not expect a double-peak in the Brillouin spectrum, and thus one could not distinguish and measure the properties of individual components in the probed focal volume [36]. Our results thus encourage a careful consideration of the spatial scales in Brillouin microscopy with respect to the biological structures of interest, especially when optimizing the relevant optical imaging parameters as well as the sample’s relative orientation.

Altogether, our results provide interesting avenues for future research, such as a more detailed and comprehensive mechanical characterization of the ECM *in-vivo*, in terms of the elasticity tensor. From a biological viewpoint, the ability to image mechanical differences inside tissues allows deciphering how tissue elasticity and viscosity contribute to development and organogenesis, where the interplay between mechanical properties, forces and signaling determines size and shape. Several major open questions exist in relation to pattern formation during tube morphogenesis, which could now be addressed in future experiments: Does the ECM have a universal role for tube elongation? How are branch points determined? How does a tissue decide when to start or stop elongating? Our results are encouraging for future work which aims to answer these questions and validate how conserved the identified mechanism are across size scales and among different tubular structures. Thus, our work opens the door to performing such delicate mechanical measurements in space and time and might shed new light on the role of ECM stiffness for tube elongation in vertebrates.

## Acknowledgements

We acknowledge K. Elsayad for fruitful discussions and initial advice regarding Brillouin microscopy. We are grateful to the EMBL EM core facility (EMCF) for support, and in particular Giulia Mizzon who performed the EM experiments‥ We thank J. Czuchnowski for help with Figure 1A. We also thank M. Bagnat and V. Mulero for kindly sharing the col9a2:CAAX-GFP, sagG214 and UAS:mCherry-NTR zebrafish lines used in this study.

## References

1. A. J. Engler, S. Sen, H. L. Sweeney, and D. E. Discher, “Matrix Elasticity Directs Stem Cell Lineage Specification,” Cell 126, 677–689 (2006).

2. D. J. Andrew and A. J. Ewald, “Morphogenesis of epithelial tubes: Insights into tube formation, elongation, and elaboration,” Dev. Biol. 341, 34–55 (2010).

3. J. Crest, A. Diz-Muñoz, D. Chen, D. A. Fletcher, and D. Bilder, “Organ sculpting by patterned extracellular matrix stiffness,” Elife 6, 1–16 (2017).

4. O. Campàs, T. Mammoto, S. Hasso, R. A. Sperling, D. O’Connell, A. G. Bischof, R. Maas, D. A. Weitz, L. Mahadevan, and D. E. Ingber, “Quantifying cell-generated mechanical forces within living embryonic tissues,” Nat. Methods 11, 183–189 (2014).

5. K. Sugimura, P.-F. Lenne, and F. Graner, “Measuring forces and stresses in situ in living tissues,” Development 143, 186–196 (2016).

6. M. Krieg, G. Fläschner, D. Alsteens, B. M. Gaub, W. H. Roos, G. J. L. Wuite, H. E. Gaub, C. Gerber, Y. F. Dufrêne, and D. J. Müller, “Atomic force microscopy-based mechanobiology,” Nat. Rev. Phys. (2018).

7. K. Guevorkian and J.-L. Maître, “Micropipette aspiration,” in (2017), pp. 187–201.

8. M. L. Gardel, J. H. Shin, F. C. MacKintosh, L. Mahadevan, P. Matsudaira, and D. A. Weitz, “Elastic behavior of cross-linked and bundled actin networks,” Science (80-.). 304, 1301–1305 (2004).

9. H. Zhang and K.-K. Liu, “Optical tweezers for single cells,” J. R. Soc. Interface 5, 671–690 (2008).

10. M. E. Dolega, M. Delarue, F. Ingremeau, J. Prost, A. Delon, and G. Cappello, “Cell-like pressure sensors reveal increase of mechanical stress towards the core of multicellular spheroids under compression,” Nat. Commun. 8, 1–9 (2017).

11. G. Scarcelli and S. H. Yun, “Confocal Brillouin microscopy for three-dimensional mechanical imaging,” Nat Phot. 2, 39–43 (2008).

12. G. Scarcelli, W. J. Polacheck, H. T. Nia, K. Patel, A. J. Grodzinsky, R. D. Kamm, and S. H. Yun, “Noncontact three-dimensional mapping of intracellular hydromechanical properties by Brillouin microscopy,” Nat. Methods 12, 1132–1134 (2015).

13. G. Antonacci and S. Braakman, “Biomechanics of subcellular structures by non-invasive Brillouin microscopy,” Sci. Rep. 6, 1–7 (2016).

14. G. Antonacci, V. de Turris, A. Rosa, and G. Ruocco, “Background-deflection Brillouin microscopy reveals altered biomechanics of intracellular stress granules by ALS protein FUS,” Commun. Biol. 1, 139 (2018).

15. G. Scarcelli, P. Kim, and S. H. Yun, “Ex-vivo measurement of age-related stiffening in the crystalline lens by Brillouin optical microscopy,” Biophys. J. 101, 1539–1545 (2011).

16. G. Scarcelli and S. H. Yun, “Ex-vivo Brillouin optical microscopy of the human eye,” Opt. Express 20, 9197 (2012).

17. G. Scarcelli, S. Besner, R. Pineda, P. Kalout, and S. H. Yun, “Ex-vivo biomechanical mapping of normal and keratoconus corneas.,” JAMA Ophthalmol. 133, 480–482 (2015).

18. R. Schlüßler, S. Möllmert, S. Abuhattum, G. Cojoc, P. Müller, K. Kim, C. Möckel, C. Zimmermann, J. Czarske, and J. Guck, “Mechanical mapping of spinal cord growth and repair in living zebrafish larvae by brillouin imaging,” Biophys. J. 115, 911–923 (2018).

19. F. Palombo, C. P. Winlove, R. S. Edginton, E. Green, N. Stone, S. Caponi, M. Madami, and D. Fioretto, “Biomechanics of fibrous proteins of the extracellular matrix studied by Brillouin scattering,” J. R. Soc. Interface 11, (2014).

20. K. Elsayad, S. Werner, M. Gallemi, J. Kong, E. R. Sanchez Guajardo, L. Zhang, Y. Jaillais, T. Greb, and Y. Belkhadir, “Mapping the subcellular mechanical properties of live cells in tissues with fluorescence emission-Brillouin imaging,” Sci. Signal. 9, rs5–rs5 (2016).

21. D. L. Stemple, “Structure and function of the notochord: an essential organ for chordate development,” Development 132, 2503–2512 (2005).

22. D. S. Adams, R. Keller, and M. A. Koehl, “The mechanics of notochord elongation, straightening and stiffening in the embryo of Xenopus laevis.,” Development 110, 115–30 (1990).

23. K. Ellis, B. D. Hoffman, and M. Bagnat, “The vacuole within,” Bioarchitecture 3, 64–68 (2013).

24. M. J. Parsons, S. M. Pollard, L. Saúde, B. Feldman, P. Coutinho, E. M. A. Hirst, and D. L. Stemple, “Zebrafish mutants identify an essential role for laminins in notochord formation.,” Development 129, 3137–46 (2002).

25. S. Grotmol, H. Kryvi, R. Keynes, C. Krossoy, K. Nordvik, and G. K. Totland, “Stepwise enforcement ofthe notochord and its intersection with the myoseptum: an evolutionary path leading to development of the vertebra?,” J. Anat. 209, 339–357 (2006).

26. J. Garcia, J. Bagwell, B. Njaine, J. Norman, D. S. Levic, S. Wopat, S. E. Miller, X. Liu, J. W. Locasale, D. Y. R. Stainier, and M. Bagnat, “Sheath Cell Invasion and Trans-differentiation Repair Mechanical Damage Caused by Loss of Caveolae in the Zebrafish Notochord,” Curr. Biol. 27, 1982–1989.e3 (2017).

27. J. M. Gansner, B. A. Mendelsohn, K. A. Hultman, S. L. Johnson, and J. D. Gitlin, “Essential role of lysyl oxidases in notochord development,” Dev. Biol. 307, 202–213 (2007).

28. F. Scarponi, S. Mattana, S. Corezzi, S. Caponi, L. Comez, P. Sassi, A. Morresi, M. Paolantoni, L. Urbanelli, C. Emiliani, L. Roscini, L. Corte, G. Cardinali, F. Palombo, J. R. Sandercock, and D. Fioretto, “High-Performance Versatile Setup for Simultaneous Brillouin-Raman Microspectroscopy,” Phys. Rev. X 7, 31015 (2017).

29. G. Scarcelli, R. Pineda, and S. H. Yun, “Brillouin Optical Microscopy for Corneal Biomechanics,” Investig. Opthalmology Vis. Sci. 53, 185 (2012).

30. G. Scarcelli and S. H. Yun, “Multistage VIPA etalons for high-extinction parallel Brillouin spectroscopy,” Opt. Express 19, 10913 (2011).

31. E. Edrei, M. C. Gather, and G. Scarcelli, “Integration of spectral coronagraphy within VIPA-based spectrometers for high extinction Brillouin imaging,” Opt. Express 25, 6895–6903 (2017).

32. M. Yamamoto, R. Morita, T. Mizoguchi, H. Matsuo, M. Isoda, T. Ishitani, A. B. Chitnis, K. Matsumoto, J. G. Crump, K. Hozumi, S. Yonemura, K. Kawakami, and M. Itoh, “Mib-Jag1-Notch signalling regulates patterning and structural roles of the notochord by controlling cell-fate decisions,” Development 137, 2527–2537 (2010).

33. J. M. Davison, C. M. Akitake, M. G. Goll, J. M. Rhee, N. Gosse, H. Baier, M. E. Halpern, S. D. Leach, and M. J. Parsons, “Transactivation from Gal4-VP16 transgenic insertions for tissue-specific cell labeling and ablation in zebrafish,” Dev. Biol. 304, 811–824 (2007).

34. S. Durdu, M. Iskar, C. Revenu, N. Schieber, A. Kunze, P. Bork, Y. Schwab, and D. Gilmour, “Luminal signalling links cell communication to tissue architecture during organogenesis,” Nature 515, 120–124 (2014).

35. N. L. Schieber, S. J. Nixon, R. I. Webb, V. M. J. Oorschot, and R. G. Parton, “Modern Approaches for Ultrastructural Analysis of the Zebrafish Embryo,” in (2010), pp. 425–442.

36. S. Mattana, “Non-contact mechanical and chemical analysis of single living cells by micro-spectroscopic techniques,” Light Sci. Appl. 7, e17139 (2018).

